# The phylogenetic and global distribution of bacterial polyhydroxyalkanoate bioplastic degrading genes

**DOI:** 10.1101/2020.05.08.085522

**Authors:** V. R. Viljakainen, L. A. Hug

## Abstract

Polyhydroxyalkanoates (PHAs) are a family of microbially-made polyesters commercialized as biodegradable plastics. PHA production rates are predicted to increase as concerns around environmental plastic contamination and limited fossil fuel resources have increased the importance of biodegradable and bio-based plastic alternatives. Microbially-produced PHA depolymerases are the key enzymes mediating PHA biodegradation, but only a few PHA depolymerases have been well-characterized and screens employing metagenomic sequence data are lacking. Here, we used 3,078 metagenomes to analyze the distribution of PHA depolymerases in microbial communities from diverse aquatic, terrestrial and waste management systems. We significantly expand the recognized diversity of this protein family by screening 1,914 Gb of sequence data and identifying 13,869 putative PHA depolymerases in 1,295 metagenomes. Our results indicate that PHA depolymerases are unevenly distributed across environments. We predicted the highest frequency of PHA depolymerases in wastewater systems and the lowest in marine and thermal springs. In tandem, we screened 5,290 metagenome-assembled genomes to describe the phylogenetic distribution of PHA depolymerases, which is substantially broader compared to current cultured representatives. The Proteobacteria and Bacteroidota are key lineages encoding PHA depolymerases, but PHA depolymerases were predicted from members of the Bdellovibrionota, Methylomirabilota, Actinobacteriota, Firmicutes, Spirochaetota, Desulfobacterota, Myxococcota and Planctomycetota.

**Originality/Significance Statement:** Biodegradable plastics like polyhydroxyalkanoates (PHAs) are a hot topic, following ubiquitous environmental plastic contamination, government bans on single-use plastics, and a growing need for sustainable alternatives to petroleum-based plastics. Understanding the microbial conversion of PHAs in the environment and finding biomolecular tools that can act on PHAs is increasingly important as PHAs grow in popularity. In this study, we screened thousands of metagenomes and metagenome-assembled genomes (MAGs) to substantially increase the recognized diversity of PHA depolymerases, the key enzymes mediating PHA biodegradation. We use datasets from seven continents to provide a global summary of the distribution of PHA depolymerase genes in natural environments and waste-management systems. In tandem, we increase the number of described phylum-level lineages with PHA biodegradation potential. This work contributes a new understanding of the phylogenetic and environmental distribution of PHA depolymerases.

## Introduction

Global concerns associated with environmental plastic contamination and fossil fuel use have led to a demand for sustainable alternatives to petroleum-based plastics^1^. Increased public awareness coupled with ongoing legislative changes aimed at reducing single-use plastics are promoting a pivot to bio-based and biodegradable alternatives like polyhydroxyalkanoates (PHAs)^2,3^. Conventional plastics rely on fossil resources and are extremely recalcitrant to degradation, causing them to persist upon disposal^4^. Annual plastic production now exceeds 380 million tons, with most plastics experiencing a short service lifespan before being discarded^5^. It is estimated that 79% of plastic waste accumulates in landfills or the environment^5^ where it can cause ecological damage^6^.

PHAs offer an exciting solution to curb environmental damage caused by liberal plastic use because they are made from renewable carbon resources (bio-based) and biodegrade into non-harmful products in natural environments through the action of microbially-secreted PHA depolymerases^7-12^. PHAs are a naturally produced family of polyesters synthesized by bacteria and archaea as intracellular energy and carbon storage compounds^13,14^. Commercial PHAs currently represent a small fraction (1.4%) of the bioplastic market^15^ but show the highest growth rates with production capacities forecasted to more than triple in the next five years^16^.

Rates of environmental plastic degradation are influenced by polymer properties (*e*.*g*., available surface area, monomeric composition, additives), the extant microbial community, and abiotic factors (*e*.*g*., temperature, UV, pH)^17^. PHA biodegradation is largely reliant on the abundance and diversity of microorganisms encoding extracellular PHA depolymerases (the key PHA degrading enzymes)^18,19^. PHAs are broadly classified as either short-chain length (PHAscl) with 3-5 carbon atoms per monomer or medium-chain length (PHAmcl) with 6-15 carbon atoms per monomer, where the size of the monomer impacts enzyme substrate specificity^13^.

PHA depolymerases are a diverse family of intracellular and extracellular carboxylesterases and are members of the alpha/beta-hydrolase fold family^20^. To date, approximately 35 PHA depolymerases have been isolated and biochemically characterized (Supp Data 1 and references within), from organisms within the phyla Ascomycetes, Firmicutes, Proteobacteria (most) and Actinobacteria. PHA depolymerases are classified based on subcellular location (extracellular or intracellular), substrate specificity (mcl [EC 3.1.1.76] or scl [EC 3.1.1.75]), and features of the catalytic domain^13,20^. Intracellular PHA (inPHA) depolymerases mobilize amorphous PHA stores in intracellular, protein-associated granules referred to as carbonosomes^21^. Extracellular PHA (ePHA) depolymerases are secreted from the cell to scavenge exogenous semi-crystalline PHAs found in the environment^13^, and are the focus of this project.

Early culture-based studies have isolated ePHAscl-degrading microorganisms from different environments including soil, compost, sewage and aquatic environments, but indicate that ePHAmcl degraders appear to be relatively rare^7,13,19^. Reflecting these findings, relatively few ePHAmcl depolymerases have been characterized compared to ePHAscl depolymerases^20,22^. All known ePHA depolymerases contain a catalytic triad composed of a serine (embedded in a lipase box motif), histidine, and aspartic acid^20^. ePHAscl depolymerases typically consist of a signal peptide, a catalytic domain, a linker domain and a C-terminal substrate binding domain^13^. There are two recognized ePHAscl depolymerase subtypes (denoted type 1 [T1] and type 2 [T2]) which differ in the arrangement of their catalytic domains. In ePHAsclT1 depolymerases, the oxyanion hole is N-terminal to the lipase box, whereas in ePHAsclT2 depolymerases the oxyanion hole is C-terminal to the lipase box^13^. Characterized ePHAmcl depolymerases do not exhibit a C-terminal substrate binding domain; instead the N-terminal region is presumed to mediate substrate binding^13^. ePHAmcl depolymerases share no significant homology with ePHAscl depolymerases except for the catalytic triad^20^ which is non-specific as it is shared by all carboxylesterases^23^.

With PHA production levels expected to increase, a strong understanding of the diversity and distribution of PHA-degrading enzymes and organisms is critical for understanding the impact of PHAs on the environment and to reveal biomolecular tools useful for processing these plastics^19^. Culture-based research has developed a foundational understanding of the enzymes and organisms mediating PHA biodegradation (reviewed by ^13^). However, many microbial lineages are currently unamenable to culturing and are only recognized from culture-independent analyses^24^. With the advent of high-throughput sequencing and metagenomics, there has been a proliferation of publicly available DNA sequence data from microbial communities inhabiting diverse global environments^25,26^, enabling broad surveys of gene distributions^27-29^.

Here, we employed a culture-independent approach and leveraged large amounts of publicly available environmental sequence data to characterize the phylogenetic and environmental distribution of PHA depolymerases and predict novel PHA-degrading enzymes. We screened 5,290 metagenome-assembled genomes (MAGs) and 3,078 metagenomes sequenced from globally distributed natural environments and waste-management systems. Biochemically characterized ePHA depolymerases were used to seed a homology-based search to identity novel PHA depolymerase candidates from different environments and across the bacterial tree of life. Our analyses significantly expand the recognized diversity of PHA depolymerases and inform our understanding of the frequency of PHA depolymerases in the environment, with implications for the microbial conversion of PHA litter. Furthermore, we identify enzymes and organisms that may prove suitable for future bioremediation, chemical processing or biotechnological applications.

## Methods

### Identification of ePHA depolymerases

A reference set of biochemically validated ePHA depolymerases were gathered from the literature (Supp Data 1) and used as queries in a BLASTp search (threshold e< 1×10^−10^) against 3,078 assembled and annotated metagenomes (Supp Data 2) and 5,290 MAGs (Supp Data 3) that were publicly available on the Integrated Microbial Genomes and Microbiomes system (IMG/M) https://img.jgi.doe.gov/m/^26^. Datasets screened were publicly available, but permissions were requested from PIs associated with the datasets. Many metagenomes came from datasets associated with publications, including^30-45^. A series of curation steps were taken to increase PHA depolymerase prediction accuracy and to reduce or eliminate non-functional homologs. Briefly, curation included eliminating severely truncated hits, ensuring key catalytic residues were present using multiple sequence alignments (MSAs), and protein-tree based curation using tree topology and branch length of hits placed among reference enzymes (for detailed curation notes see Supp Data 4). BLAST output data for the reference query sequence with the lowest e-value and a summary of the average, minimum, and maximum bitscores, identities, e-values, and lengths of predicted PHA depolymerases can be found in Supp Data 5.Signal peptides were predicted for predicted PHA depolymerases using Signal P 5.0 (https://services.healthtech.dtu.dk/service.php?SignalP) with default settings using both gram negative and gram positive predictions^46^.

### Dataset selection, ecosystem classification & normalization

All 3,078 metagenomic datasets had a minimum of 0.5 Mb of assembled sequence data and were selected to analyze the environmental distribution of PHA depolymerases. All metagenomic datasets had environmental metadata available in accordance with the GOLD five-level ecosystem classification system ([1] ecosystem [2] ecosystem category [3] ecosystem type [4] ecosystem subtype and [5] specific ecosystem)^47^. All environmental data provided was used without modification with the exception of 8 metagenomes with ecosystem subtype classifications of loam (3) and peat (5) which were modified to soil to match other loam and peat samples. Additionally, sediment samples were inconsistently classified at the [5] specific ecosystem type or [3] ecosystem type levels, which was remedied by manually grouping all aquatic sediment samples together as an ecosystem type (see Supp Data 2 for modifications).

Collectively, 104 unique GOLD ecosystems were described (Supp Data 2). Less than two percent of metagenomes (40/3,078) lacked latitude and longitude locations, which were input manually using data given in an associated study, or approximate geocoordinates were used given samples’ described geographical locations (Supp Data 2). To account for differing dataset sizes, we calculated the frequency (hits/Gb) of PHA depolymerases occurring in metagenomes and MAGs (Supp Fig 1; Supp Data 3). Frequencies for different GOLD level ecosystem classifications were derived by dividing the total number of enzymes identified in metagenomes from an ecosystem category or subtype by the sum total of the sequence data screened from metagenomes occurring within a group. Frequencies in the MAGs for different phyla were calculated by dividing the total number of enzymes identified from a phylum by the sum total of the sequence data screened from MAGs of that phylum. Gene frequencies were calculated in this manner to normalize based on clade size.

**Figure 1:**
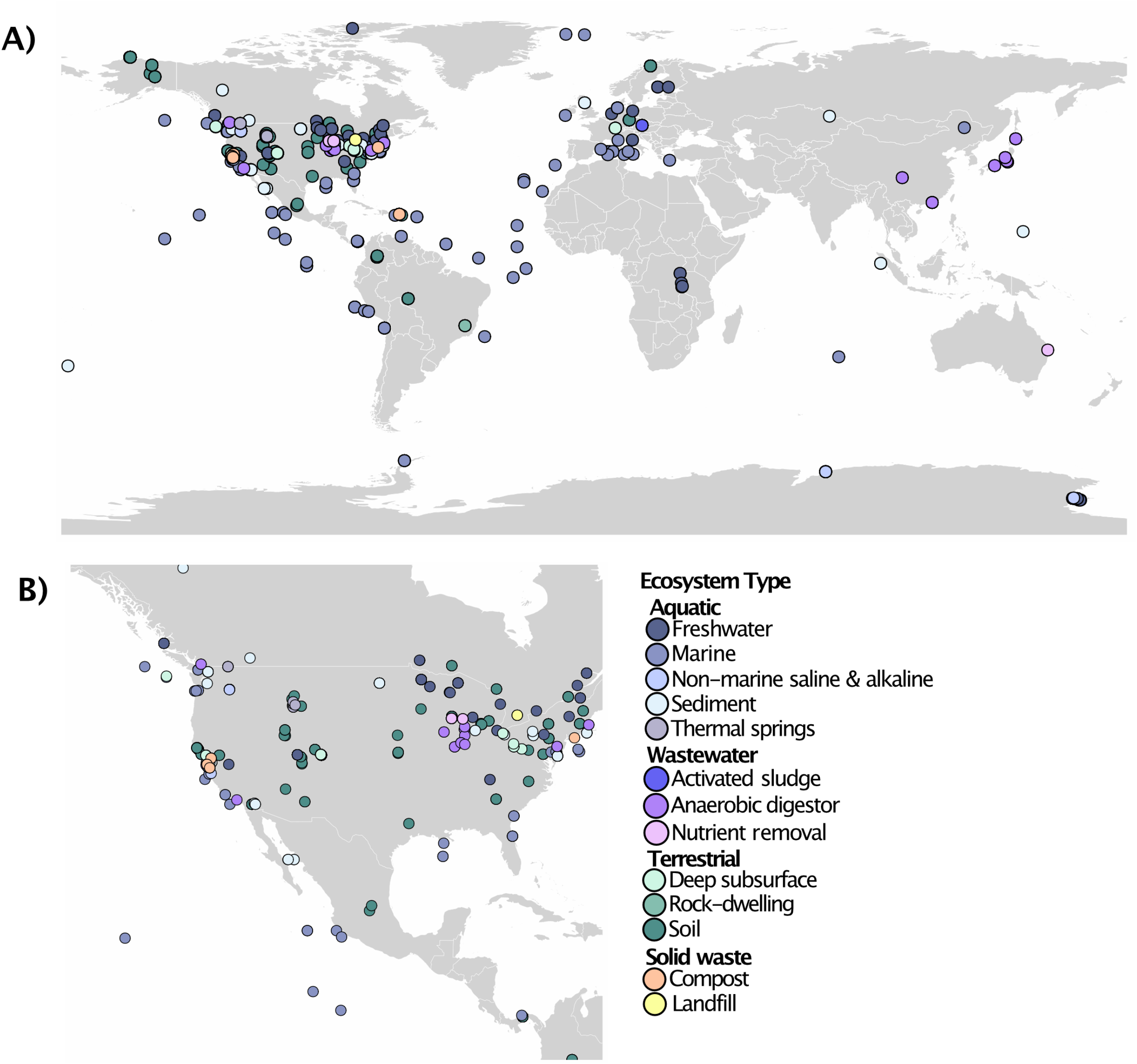
A total of 13,869 PHA depolymerases were predicted in 1,295 metagenomes from aquatic (417), terrestrial (764), solid waste (23) and wastewater (91) environments. **A)** World map showing geocoordinates of samples with metagenomes containing one or more predicted PHA depolymerase, colored by GOLD ecosystem type. Metagenomes sequenced from all seven continents were included in our dataset but were unequally represented, with certain regions having more sampled locations and sample types than others. **B)** Metagenomes from North America were the most heavily represented in our dataset and nearly every ecosystem type showed some evidence for PHA depolymerase genes within this region. Several samples share geocoordinates and represent samples taken at different depths, samples taken within close proximity to each other, or at different time points.

### Taxonomic affiliation of PHA depolymerases

Phylogenetic affiliations of PHA depolymerases were predicted for the unbinned PHA depolymerase-encoding scaffolds with a minimum length of 2,000 bps (7,260 scaffolds) using a diamond-blast search^48^ against NCBI’s non-redundant database and subsequently conducting a lowest common ancestor analysis (LCA) with MEGAN6 v6.19.5^49^ using GTDB taxonomy^50^. For MAGs, 143 MAGs encoding predicted PHA depolymerases were assessed using CheckM v1.0.13^51^, with MAGs kept for on-going analyses if they scored >50% complete and <10% contaminated (Supp Data 3). MAGs were subsequently dereplicated using dRep v. 2.5.3^52^. GTDB-tk^50^, as implemented on Kbase^53^, was used to taxonomically identify dereplicated MAGs. Finally, a concatenated ribosomal protein (rp) tree was constructed to phylogenetically place the dereplicated MAGs that were >50% complete and <10% redundant (Supp Data 3). A dataset of 16 rps (rpL2, L3, L4, L5, L6, L14, L15, L16, L18, L22, L24, S3, S8, S10, S17 and S19) were gathered from MAGs and independently aligned. Alignments were masked to remove columns comprised of 95%+ gaps. MAGs with <50% of expected ungapped residues over the concatenated alignment were excluded. Phyla containing predicted PHA depolymerase-encoding MAGs were selected from a bacterial reference set^24^ and included in the final concatenated alignment. The final alignment was composed of 2,789 positions, 886 reference taxa and 98 MAGs. The final concatenated rp tree was inferred using FastTree^54^ as implemented in Geneious (https://www.geneious.com) and the tree was visualized using Itol (https://itol.embl.de/)^55^.

## Results

### Environmental distribution of predicted PHA depolymerases

All IMG/M metagenomes included in this study had a minimum of 0.5 Mb of assembled sequence data and had available GOLD environmental metadata (Supp Data 2). The 3,078 metagenomes were classified into 104 specific ecosystems from the GOLD hierarchy level 3 (Supp Data 2). Metagenomes were treated at the level of ecosystem category (level 1: aquatic, terrestrial, solid waste or wastewater) and ecosystem type (level 2: 14 subcategories). We focus on specific ecosystems or metagenomes only for select notable cases, however all data including raw number of predicted PHA depolymerases, presence/absence, and frequency of predicted enzymes identified from each metagenome and each specific ecosystem type can be found in Supp Data 2.

A total of 13,869 PHA depolymerases (13,377 unique) were predicted in 1,295 metagenomes from aquatic (417/1,492 = 27.9%), solid waste (23/34 =67.6%), terrestrial (764/1,323 = 57.7%) and wastewater (91/229 = 39.7%) environments (Fig 1). Of these, 43% (5,949/13,869) had predicted signal peptides. Most metagenomes had a low frequency of predicted PHA depolymerases (0.3-10 hits/Gb) (Supp Fig 1, Supp Data 2). Looking broadly at different ecosystem categories, the highest frequency of predicted PHA depolymerases occurred in wastewater systems (10.73 hits/Gb) and terrestrial environments (8.22 hits/Gb) whereas solid waste and aquatic systems had averages of 5.53 and 4.09 hits/Gb respectively (Table 1, Supp Data 2).

**Table 1:** Environmental distribution of 13,869 predicted PHA depolymerases (222 ePHAmcl, 10,245 ePHAsclT1 and 3,402 ePHAsclT2 depolymerases) identified by screening 3,078 metagenomes. Number of metagenomes screened, percent of metagenomes with at least one or more predicted PHA depolymerase, the number of predicted PHA depolymerases identified, and the frequency of predicted PHA depolymerases (hits/Gb) per ecosystem type are included.

| Metagenomes Screened (n) | # of predicted PHA depolymerases |  |  |  | % of metagenomes with predicted PHA depolymerases |  |  |  | Frequency of predicted PHA depolymerases (hits/Gb) |  |  |  |
| --- | --- | --- | --- | --- | --- | --- | --- | --- | --- | --- | --- | --- |
| Ecosystem | sclT1 | sclT2 | mcl | Total | sclT1 | sclT2 | mcl | Total | sclT1 | sclT2 | mcl | Total |
| **Aquatic (1,492)** |  |  |  |  |  |  |  |  |  |  |  |  |
| Freshwater (481) | 648 | 140 | 23 | 811 | 26.0 | 13.5 | 3.3 | 27.4 | 3.94 | 0.85 | 0.14 | 4.93 |
| Marine (373) | 187 | 126 | 20 | 333 | 19.3 | 21.4 | 4 | 31.4 | 1.06 | 0.71 | 0.11 | 1.88 |
| Saline and Alkaline (75) | 41 | 18 | 0 | 59 | 24.0 | 20.0 | 0 | 34.7 | 2.91 | 1.28 | 0.00 | 4.18 |
| Sediment (442) | 488 | 286 | 10 | 784 | 26.2 | 23.3 | 2.3 | 30.5 | 3.76 | 2.21 | 0.08 | 6.05 |
| Thermal springs (121) | 15 | 1 | 0 | 16 | 5.8 | 0.8 | 0 | 5.8 | 3.14 | 0.21 | 0.00 | 3.34 |
|  | 1,379 | 571 | 53 | 2,003 | 22.7 | 17.7 | 2.7 | 27.9 | 2.81 | 1.16 | 0.11 | 4.09 |
| **Solid Waste (34)** |  |  |  |  |  |  |  |  |  |  |  |  |
| Compost (30) | 27 | 29 | 0 | 56 | 33.3 | 63.3 | 0 | 66.7 | 4.52 | 4.85 | 0.00 | 9.37 |
| Landfill (4) | 19 | 9 | 0 | 28 | 75.0 | 75.0 | 0 | 75 | 2.06 | 0.98 | 0.00 | 3.04 |
|  | 46 | 38 | 0 | 84 | 38.2 | 64.7 | 0 | 67.6 | 1.78 | 1.91 | 0.00 | 5.53 |
| **Terrestrial (1,323)** |  |  |  |  |  |  |  |  |  |  |  |  |
| Deep subsurface (104) | 59 | 16 | 1 | 76 | 18.3 | 7.7 | 1 | 20.2 | 6.19 | 1.68 | 0.10 | 7.97 |
| Rock-dwelling (3) | 28 | 5 | 0 | 33 | 100 | 66.7 | 0 | 100 | 9.17 | 1.64 | 0.00 | 10.81 |
| Soil (1,211) | 8,331 | 2,374 | 130 | 10,835 | 57.1 | 43.8 | 7.7 | 61.1 | 6.32 | 1.80 | 0.10 | 8.22 |
| Volcanic (5) | 0 | 0 | 0 | 0 | 0 | 0 | 0 | 0 | 0.00 | 0.00 | 0.00 | 0.00 |
|  | 8,418 | 2,395 | 131 | 10,944 | 53.9 | 40.8 | 7.1 | 57.7 | 6.33 | 1.80 | 0.10 | 8.22 |
| **Wastewater (229)** |  |  |  |  |  |  |  |  |  |  |  |  |
| Activated Sludge (140) | 133 | 73 | 11 | 217 | 9.3 | 5.7 | 5 | 9.3 | 8.47 | 4.65 | 0.70 | 13.82 |
| Anaerobic digester (54) | 152 | 179 | 7 | 338 | 70.4 | 87 | 13 | 94.4 | 3.37 | 3.97 | 0.16 | 7.50 |
| Nutrient removal (35) | 117 | 146 | 20 | 283 | 60 | 77.1 | 11.4 | 77.1 | 6.76 | 8.43 | 1.15 | 16.34 |
|  | 402 | 398 | 38 | 838 | 31.4 | 35.8 | 7.9 | 39.7 | 5.15 | 5.10 | 0.49 | 10.73 |
| **Total: 3,078** | **10,245** | **3,402** | **222** | **13,869** | **36.9** | **29.5** | **5** | **42.1** | **5.35** | **1.78** | **0.12** | **7.24** |

1. **Aquatic:** across all aquatic ecosystem types, around 30% (±5%) of metagenomes had predicted PHA depolymerases, with the exception of thermal spring metagenomes (7/121= 5.8%) (Table 1). Despite this finding, marine ecosystem metagenomes had the lowest frequency of predicted enzymes within aquatic environments (1.88 hits/Gb). Notably, no metagenomes from marine samples nor thermal springs contained any predicted PHAmcl enzymes. The highest frequency of predicted PHA depolymerases in this category occurred in aquatic sediments (6.05 hits/Gb). (Table 1).
2. **Solid waste:** 66.7% (20/30) of compost metagenomes and 75.0% (3/4) of landfill metagenomes had predicted PHAscl depolymerases at a frequency of 9.37 hits/Gb and 3.04 hits/Gb respectively (Table 1).
3. **Terrestrial:** Soil microbial communities have been extremely well-sampled relative to other environments. From our screen, 61.1% (740/1,211) of soil metagenomes had predicted PHA depolymerases at a frequency of 8.22 hits /Gb. Other terrestrial environments with predicted depolymerases were deep subsurface environments (21/104= 20.2%, 7.97 hits/Gb) and rock-dwelling communities (3/3 = 100.0%, 10.81 hits/Gb) (Table 1).
4. **Wastewater:** Within wastewater environments, the majority of anaerobic digestors (51/54 = 94.4%) and nutrient removal communities (27/35=77.1%) had predicted PHA depolymerases, whereas only a small fraction of activated sludge communities had any (13/140=9.3%). Despite this, activated sludge communities had a high frequency of predicted PHA depolymerases (13.82 hits/Gb), second to nutrient removal systems (16.34 hits/Gb), while anaerobic digestors were more similar to sediments and deep subsurface environments with 7.50 hits/Gb (Table 1; Supp Data 2).

### The environmental distribution of PHA depolymerase subtypes

1. **PHAmcl depolymerases**: PHAmcl depolymerases were sparsely predicted across environments, with only 222 PHAmcl depolymerases identified from 153 metagenomes (Supp Fig 2a) from aquatic (41/1,492 = 2.7%, 0.11 hits/Gb), terrestrial (94/1323 = 7.1%, 0.10 hits/Gb), and wastewater (18/229 = 7.9%, 0.49 hits/Gb) environments (Table 1). The highest frequency of PHAmcl depolymerases was detected in nutrient removal (1.15 hits/Gb) and activated sludge (0.70 hits/Gb) wastewater systems. Solid waste systems had no predicted PHAmcl depolymerases (Table 1). Of the 222 PHAmcl depolymerases predicted in this study, 208 placed in maximum-likelihood trees with reference enzymes isolated from species of *Bdellovibrio*^56^ and *Pseudomonas*^57,58^. Another 14 predicted PHAmcl depolymerases formed a second clade with enzymes recently isolated from *Streptomyces* species, which share limited sequence homology with other ePHAmcl biomarker enzymes^22,59^ (Supp Fig 2a).

**Figure 2:**
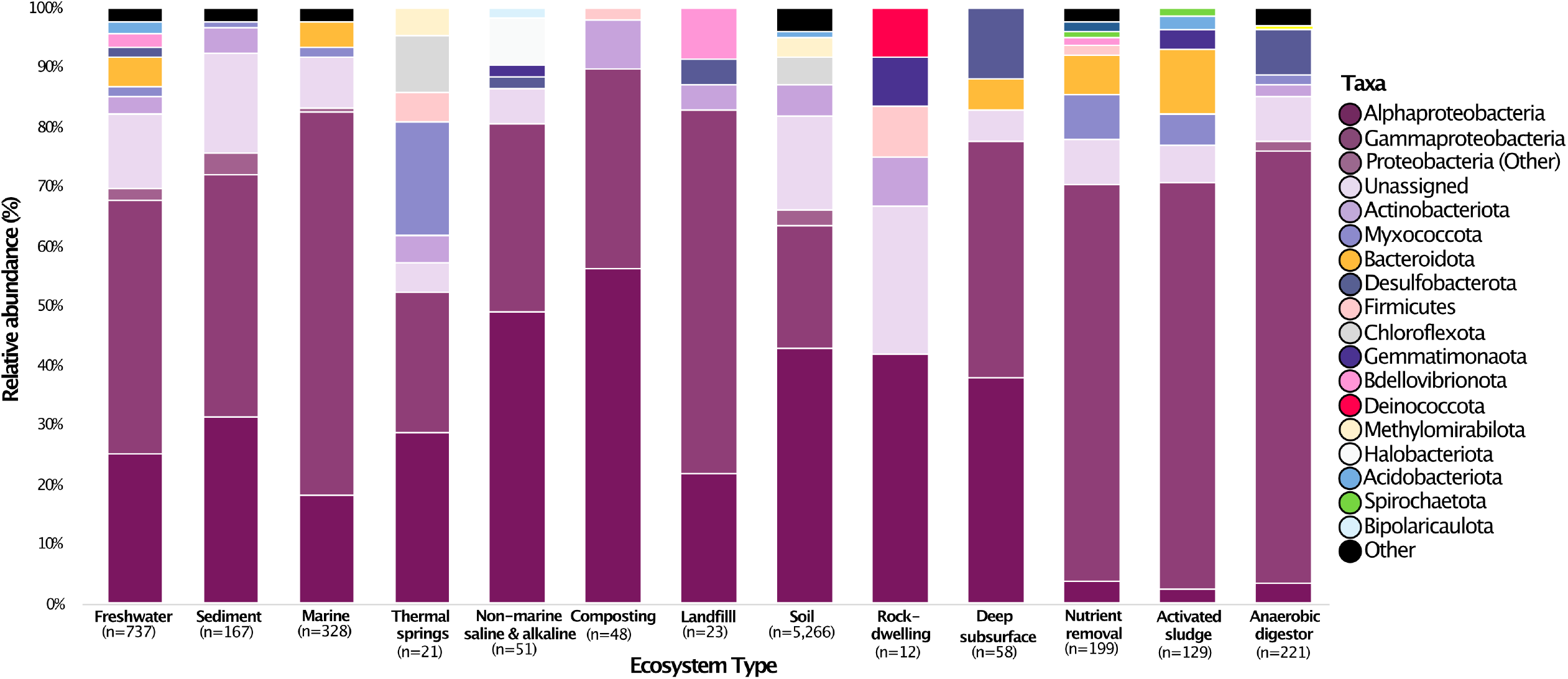
Phylogenetic affiliation of scaffolds >2000 bps in length encoding at least one predicted PHA depolymerase across different ecosystems, presented as relative abundances. “Unassigned” proteins could not be not classified at the phylum-level. Phyla that contributed less than 1% of proteins (per ecosystem type) were combined into the “other” category. Number of PHA depolymerase-encoding scaffolds for each environmental category is listed (n).
2. **PHAscl depolymerases:** Predicted PHAscl depolymerases were more prevalent across environments. A total of 10,245 PHAsclT1 depolymerases were predicted from 1,136 metagenomes (Supp Fig 2b), from aquatic (338/1,492 =22.7%, 2.81 hits/Gb), solid waste (13/34= 38.2 %, 3.03 hits/Gb), terrestrial (713 /1,323 = 53.9%, 6.33 hits/Gb), and wastewater (72/229 = 31.4%, 5.15 hits/Gb) environments (Table 1). A large number of T1 depolymerases were predicted in soil metagenomes (8,331 enzymes), but when scaled for dataset size this normalized to a frequency of 6.32 hits/Gb. All three metagenomes sequenced from rock-dwelling microbial communities encoded T1 enzymes and collectively had the highest frequency across ecosystem types with 9.17 hits/Gb. Activated sludge wastewater systems also had a higher frequency of predicted T1 enzymes with 8.47 hits/Gb.

For PHAsclT2 depolymerases, 3,402 were predicted from 908 metagenomes (Supp Fig 2c) from aquatic (264/1,492 =17.7%, 1.16 hits/Gb), solid waste (22/34 = 64.7%, 2.50 hits/Gb), terrestrial (540/1,323 = 40.8%, 1.80 hits/Gb), and wastewater (82/229 = 35.8%, 5.10 hits/Gb) environments (Table 1). Here too, a large number of hits (2,374) came from soil metagenomes, but soil did not have a particularly high frequency of enzymes (1.80 hits/Gb) compared to other ecosystem types. In fact, the frequency of predicted T2 enzymes was low (<2.5 hits/Gb) across most ecosystems with the exceptions of compost (4.85 hits/Gb) and wastewater systems (5.10 hits/Gb).

Tree files, MSAs and fasta files for all predicted PHA depolymerases and information linking hits to associated metagenome information including environmental metadata, size, scaffold read depth etc., can be found in Supp Data 2 and https://github.com/vrviljak/-The-phylogenetic-and-global-distribution-of-polyhydroxyalkanoate-bioplastic-degrading-genes.git.

### Taxonomic affiliation of unbinned PHA depolymerase-encoding scaffolds

Only a few biochemically characterized ePHA depolymerases exist, which originate from the Proteobacteria, Actinobacteriota, Firmicutes, Bdellovibrionota, and Ascomycetes (Supp Data 1). To assess the taxonomic distribution of unbinned PHA depolymerase-encoding scaffolds >2000 bp in length (7,250 scaffolds), taxonomic affiliations were inferred using MEGAN^49^ with GTDB-tk^50^ taxonomy (Supp Data 2). Most sequences were assigned to the Proteobacteria, Actinobacteriota, Chloroflexota, Methylmirabilota, Bacteroidota, and Myxococcota (Supp Fig 3). Predicted PHAmcl depolymerase-encoding scaffolds were mainly affiliated with the Myxococcota (26.0%), Proteobacteria (17.8% Gammaproteobacteria and 1.4% Alphaproteobacteria), Actinobacteriota (16.4%), and Bdellovibrionota (4.1%). Scaffolds encoding predicted PHAmcl depolymerases also originated from the Deinococcota (1), Dormibacterota (1), and Thermoplasmatota (1) (Supp Data 2). 27.4% were unclassified.

**Figure 3:**
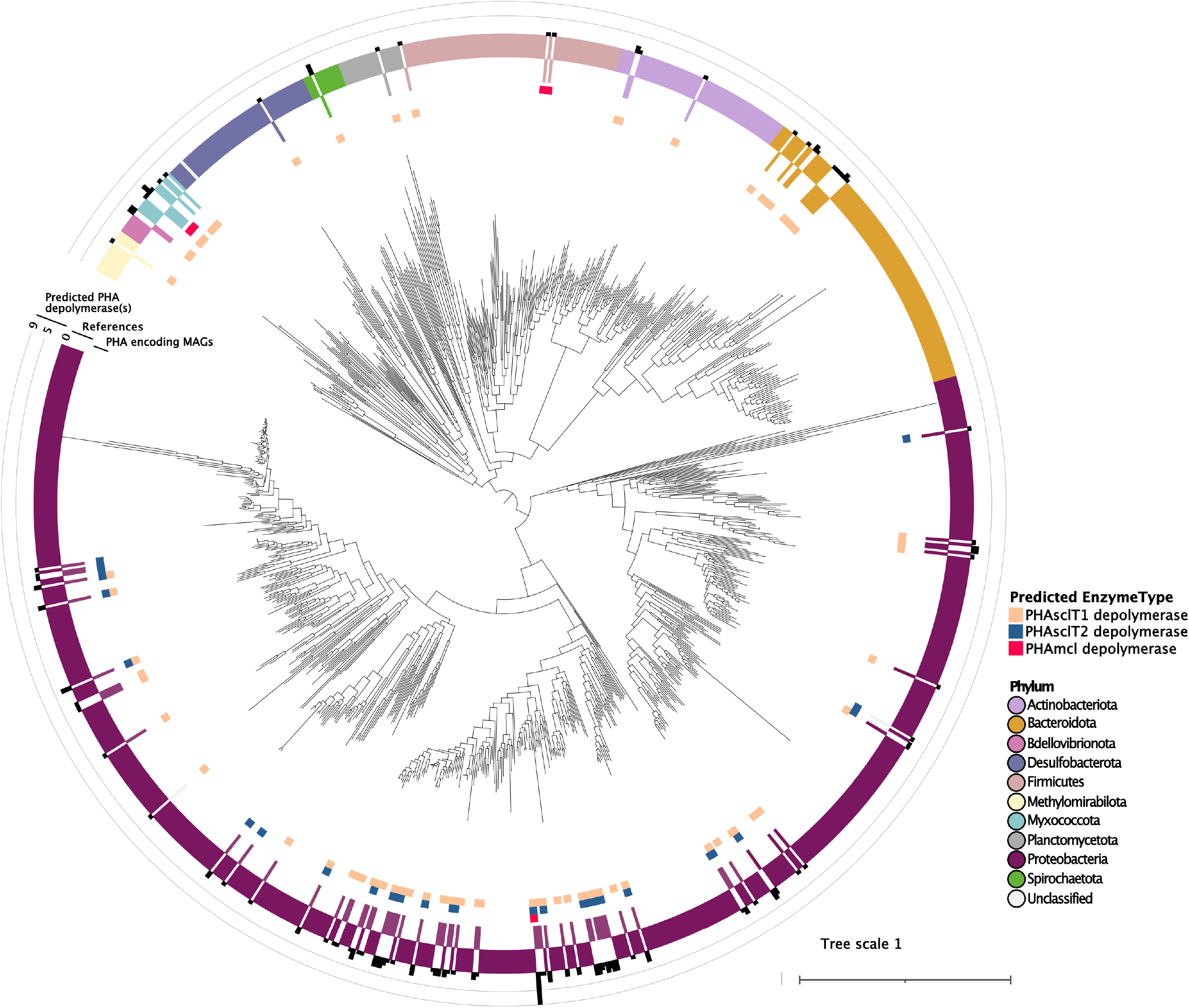
The predicted phylogenetic distribution of MAGs encoding predicted PHA depolymerases. A concatenated rp tree inferred using FastTree shows the placement of 98 high-quality, dereplicated MAGs encoding predicted PHA depolymerases placed among reference sequences selected from the bacterial domain. Tree has been rooted between the Proteobacteria and the other phyla to enhance readability. Tree scale bar indicates number of substitutions per site. The colored ring labelled “PHA-encoding MAGs” is colored to indicate the phylum-level classification for each MAG as predicted by GTDB-tk. Above this, the “References” colored ring indicates the phylum-level classification of the bacterial reference sequences. The outer ring bar plot shows the number of PHA depolymerases predicted in each MAG. The type(s) of predicted PHA depolymerase(s) (mcl, sclT1, or sclT2) found in that MAG are indicated with colored squares on the innermost rings.

The majority of predicted PHAsclT1 enzymes were on scaffolds affiliated with the Proteobacteria (64.1%), with some representation from the Actinobacteriota (5.0%), Chloroflexota (4.7%), Methylomirabilota (3.4%), Bacteroidota (2.3%), Myxococcota (1.4%) and Acidobacteriota (1.0%) (Supp Fig 3). Unclassified scaffolds represented a further 14.6%, and 17 other phyla were represented at <1% of sequences (Supp Data 2). PHAsclT2 enzymes were mostly predicted to be from the Proteobacteria (80.4%), with some in the Actinobacteriota (2.3%) (Supp Fig 3). T2 enzymes were predicted at <1% in 12 phyla, with 13.3% unclassified at the phylum level (Supp Data 2). Six PHA depolymerase scaffolds were assigned to archaeal phyla including the Halobacteriota (5), encoding predicted PHAsclT1 depolymerases, and one Thermoplasmatota scaffold encoding a predicted PHAmcl depolymerase. 5,122 predicted PHA depolymerase encoding scaffolds were classified down to the family level (2,138 or 29.5% of the phylum-level classified scaffolds were unclassified at the family-level). Family classified scaffolds were affiliated with 184 different families, including the Xanthobacteraceae (16.0%), Burkholderiaceae (13.1%), Beijerinckiaceae (4.7%), Rhodocyclaceae (3.0%), Alteromonadaceae (1.7%), Methyloligellaceae (1.6%) of the Proteobacteria, the Ktedonobacteraceae (3.09%) of the Chloroflexota, and the GWA2-73-35 (2.4%) of the Methylomirabilota. A complete family-level overview of predicted PHA depolymerases can be found in Supp Data 2.

Across ecosystem types the Proteobacteria (Alpha and Gamma) were the most abundant predicted PHA depolymerase hosts (Fig 2). Soil environments harboured the highest diversity of predicted PHA depolymerase hosts with 24 phyla predicted. Notably, several phyla were found only in soil (at <1% abundance) - the Desulfobacterota, Dormibacterota, Elusimicrobiota, Nitrospirota, Patescibacteria, Poribacteria, and Thermoplasmatota (Supp Data 2). The Myxococcota-encoded PHA depolymerases appeared in relatively high levels in thermal springs (19.1% of taxonomically assigned PHA depolymerase-encoding scaffolds) and lower levels in some other ecosystem types (1.9% in freshwater, 1.2% in sediment, 1.5% in marine, 7.5% in nutrient removal, 5.4% in activated sludge and 1.8% in anaerobic digestors). Chloroflexota-associated PHA depolymerases were also relatively abundant in thermal springs (9.5%) and only otherwise found in soil (4.5%). Five phyla were detected in all wastewater ecosystem types - the Proteobacteria, Bacteroidota, Gemmatimonadota, Myxococcota, and Spirochaetota.

Predicted PHA depolymerases co-occurred in pairs on the same assembled scaffold 39 times (1.1%), including 28 pairs of predicted PHAsclT1 depolymerases predicted to be from the Proteobacteria, Gemmatimonadota, Chloroflexota, Actinobacteriota, Myxococcota, and Bacteroidota, six pairs of predicted PHAsclT2 depolymerases from the Proteobacteria and Eremiobacterota, and one pair of predicted PHAmcl depolymerases from *Sandaracinus amylolyticus* of the Myxococcota. Predicted PHAsclT1 and PHAsclT2 depolymerases co-occurred in four cases, with three predicted as from the Gammaproteobacteria.

### Phylogenetic distribution of predicted PHA depolymerase-encoding MAGs

To further describe the phylogenetic distribution of PHA depolymerases, we separately screened 5,290 MAGs available on IMG/M for PHA depolymerases. A total of 4,927 bacterial and 363 archaeal MAGs across over 70 phyla were included in our screen, which were reconstructed from metagenomes originating from engineered (443), environmental (4,659), host-associated (186), and unclassified (2) systems (Supp Data 3). We identified a total of 143 MAGs (2.7%) encoding a total of 231 predicted PHA depolymerases in MAGs from 10 different bacterial phylum-level lineages. Of these, 71% (163/231) had predicted signal peptides. In our screen, the highest frequency of predicted PHA depolymerases (hits/Gb) were identified in the Proteobacteria (90.6 hits/Gb), followed by the Bdellovibrionota (52.5 hits/Gb), Myxococcota (47.17 hits/Gb), Methylomirabilota (28.1 hits/Gb), Actinobacteriota (20.9 hits/Gb), Bacteroidota (16.5 hits /Gb), Spirochaetota (16.1 hits /Gb), Firmicutes (12.8 hits /Gb), Desulfobacterota (5.8 hits /Gb), and Planctomycetota (3.4 hits/Gb). The remaining phyla represented in the MAG dataset had no predicted hits (Supp Data 3). 110 dereplicated PHA depolymerase-encoding MAGs were >50% complete and <10% redundant, and 98/110 encoded sufficient marker genes for inclusion in a rp-based phylogenetic tree (Fig 3). GTDB-tk successfully classified all but 5/110 MAGs (Supp Data 3), with those 5 placing among the Proteobacteria in the concatenated rp tree (Fig 3). Notably, we did not detect any MAGs with predicted PHA depolymerases from MAGs associated with the Candidate Phyla Radiation/Patescibacteria.

1. **Proteobacteria:** The vast majority of PHA depolymerases were predicted in dereplicated MAGs classified as Alphaproteobacteria (14/110) and Gammaproteobacteria (63/110). From the Alphaproteobacteria we identified MAGs from 10 families including the Hyphomicrobiaceae (3 MAGs) and the Sphingomonadaceae (4 MAGs). From the Gammaproteobacteria we identified MAGs from 15 families with most from the Burkholderiaceae (31 MAGs), the Rhodocyclaceae (10 MAGs), and the Alteromonadaceae (5 MAGs).Within the Proteobacteria, all three ePHA depolymerase types were detected (mcl = 4; sclT1 = 83; sclT2 = 45).
2. **Bacteroidota:** Beyond the Proteobacteria, 11 MAGs from the Bacteroidota encoded one (9 MAGs) or two (2 MAGs) predicted ePHAsclT1 depolymerases from seven families. An 2-12-FULL-35-15 and a GWF2-33-38 MAG each encoded two predicted ePHAsclT1 enzymes that shared high sequence similarity. Bacteroidota with one predicted PHAsclT1 depolymerase included members of the b-17BO (1), GWA2-32-17 (1), GWF2-33-38 (3), GWF2-35-48 (2), Saprospiraceae (1), and UBA2798 (1).
3. **Myxococcota:** Seven Myxococcota (formally the deltaproteobacteria) encoded predicted PHA depolymerases. Three PHAmcl depolymerases were predicted from two SG8-38 MAGs and a Polyangiaceae. A PHAsclT1 depolymerase was predicted from a Haliangiaceae, Nannocystaceae, SG8-38 and three from a Polyangiaceae MAG.
4. **Actinobacteriota:** Five Actinobacteriota MAGs from the Cellulomonadaceae, Pseudonocardiaceae and QHCA01 were predicted to encode ePHAsclT1 depolymerase(s). Three were Pseudonocardiaceae, two of which were predicted to encode two ePHAsclT1 depolymerase.
5. **Firmicutes:** Two Clostridia (Firmicutes A) both from the Ruminococcaceae family sequenced from an anerobic bioreactor were predicted to have a PHAmcl depolymerase. One UBA5603 (Firmicutes) was predicted to encode a PHAsclT1 depolymerase.
6. **Others:** Three Bacteriovoracales from the Bdellovibrionota encoded one (1 MAG) or two (2 MAGs) predicted PHAsclT1 depolymerases. A Spirochaetota from freshwater encoded three predicted PHAsclT1 depolymerases, a Desulfobacterota, a Planctomycetota and a Methylomirabilota (formally the Candidate phylum Rokubacteria) all encoded one. For a detailed overview of the PHA depolymerase complement predicted for each MAG and taxonomic affiliations, see Supp Data 3.

### MAGs encoding multiple predicted PHA depolymerases

Of the 110 dereplicated MAGs with predicted PHA depolymerases, 68 were predicted to encode one PHA depolymerase, 25 were predicted to encode two, 16 were predicted to encode three, and one, a *Herminiimonas* species (98.48% complete, 4.69% redundant) from the Burkholderiaceae family, was predicted to encode nine PHA depolymerases (Supp Data 3). The *Herminiimonas* MAG was predicted to have two closely related ePHAsclT1, four ePHAsclT2 and three closely related ePHAmcl depolymerases. MAGs from the Burkholderiaceae (6), Rhodocyclaceae (6) and Halomonadaceae (1) from the Proteobacteria all encoded three predicted PHAscl enzymes each encoding both T1 and T2 enzyme(s). One unclassified Proteobacteria that clustered closely with a Bradyrhizobiaceae reference sequence in the rp tree encoded three closely related PHAsclT1 enzymes. MAGs from the Polyangiaceae of the Myxococcota (1) and GWB1-36-13 of the Spirochaetota (1) each encoded three predicted PHAscl T1 depolymerases. Twelve of the Proteobacteria with three predicted depolymerases were from the Burkholderiales and one from the Pseudomonadales, each encoding both T1 and T2 enzyme(s).

### Phylogenetic distribution of MAG-encoded PHA depolymerase subtypes

1. **ePHAmcl depolymerases:** We predicted 8 unique ePHAmcl depolymerase genes from 6 dereplicated MAGs (Fig 4a). All PHAmcl depolymerases occurred as singletons with the exception of the three predicted in the *Herminiimonas-*related MAG, which co-occurred with six PHAscl depolymerases. Three PHAmcl depolymerases were predicted in members of the Myxococcota and two were predicted in Firmicutes. Based on ML analyses, the predicted Firmicutes ePHAmcl depolymerases were more closely related to the biochemically validated *Streptomyces* ePHAmcl depolymerases, whereas the other predicted PHAmcl enzymes were more closely related to those reference ePHA depolymerases isolated from *Pseudomonas* and *Bdellovibrio* (Fig 4a).

**Figure 4:**
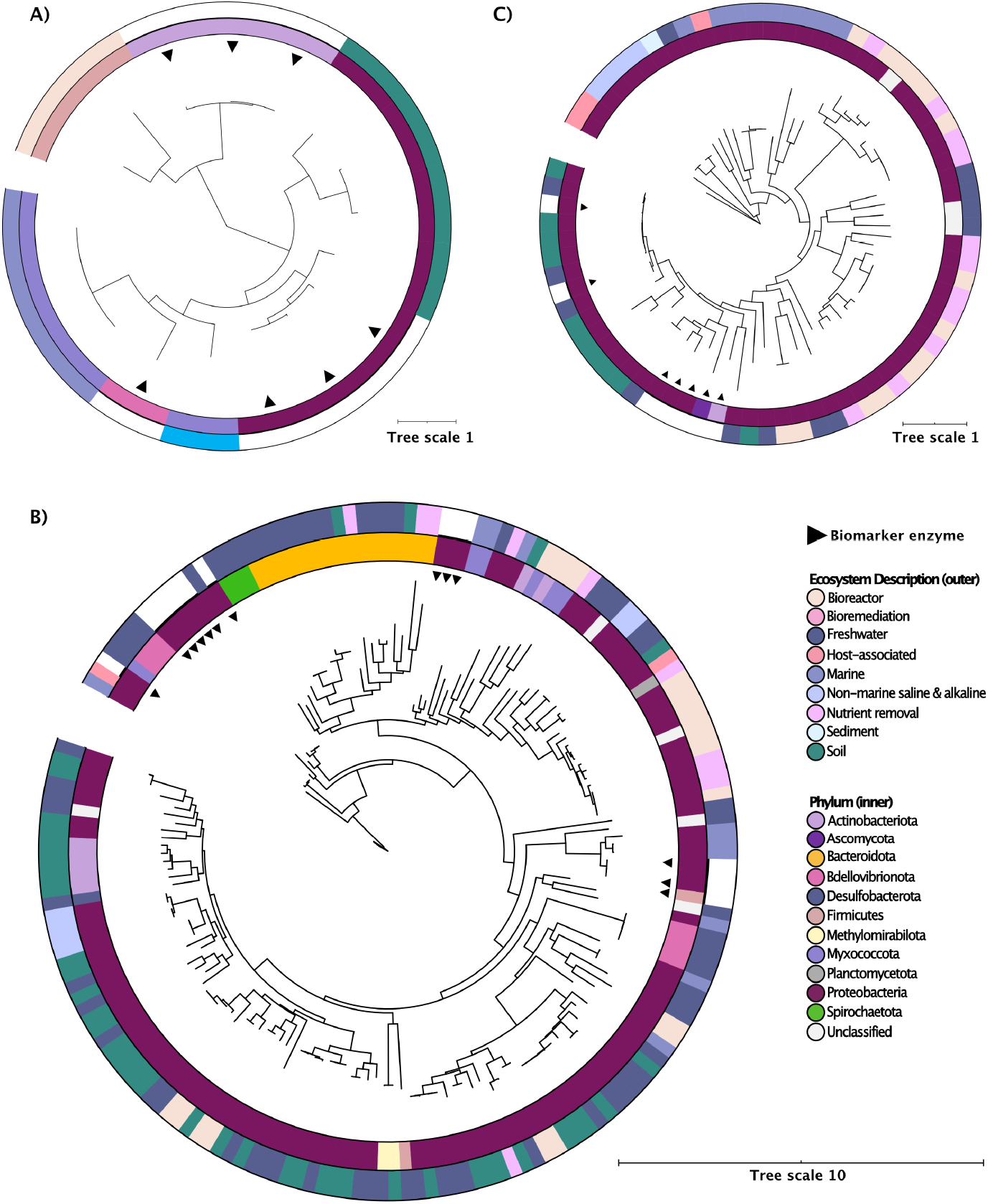
Environmental origin and phylogenetic relationships of the 231 PHA depolymerases predicted from all 143 MAGs. Tree scale bar indicates number of substitutions per site. Outer rings are colored by environmental origin of predicted PHA depolymerases. Inner rings are colored by phylum of the MAG encoding the predicted PHA depolymerase. ML trees showing **A)** 10 (8 unique) PHAmcl depolymerases **B)** 156 (133 unique) PHAscl T1 depolymerases and **C)** 65 (56 unique) PHAscl T2 depolymerases.
2. **ePHAscl depolymerases:** In our MAG screen, we predicted 109 unique ePHAsclT1 depolymerases from 84 dereplicated MAGs from the Proteobacteria (59), Bacteroidota (11), Myxococcota (4), Actinobacteriota (3), Bdellovibrionota (2) Desulfobacterota (1), Firmicutes (1), Methylomirabilota (1), Planctomycetota (1) and Spirochaetota (1) (Fig 4b). Predicted T1 enzymes occurred as mostly as singletons (62 MAGs) but also in pairs (19 MAGs), or triplets (3 MAGs). T1 and T2 enzymes were predicted to co-occur in 17 Proteobacteria MAGs. We predicted 44 unique ePHAsclT2 depolymerases in 33 dereplicated MAGs exclusively from members of the Proteobacteria (Fig 4c). Three MAGs were not classified with GTDB-tk, but placed among the Proteobacteria in the concatenated rp tree near Burkholderiales (1 MAG) and Rhizobiales (2 MAGs) reference sequences. Most predicted T2 enzymes occurred as singletons (24) or co-occurred with T1 enzymes (23). Notably, seven Burkholderiales from the Rhodocyclaceae family encoded two predicted T2 enzymes as well as a predicted T1 enzyme.

## Discussion

PHA depolymerases degrade PHA substrates in the environment and have potential applications for bioremediation, composting, and chemical recycling. The objective of this study was to increase the recognized diversity of PHA depolymerases and to elucidate their phylogenetic affiliations and occurrence in the environment. We used a homology-based search seeded with biochemically validated ePHA depolymerases followed by manual curation to identify 13,869 predicted PHA depolymerases in 1,295 annotated and assembled metagenomes from diverse global environments (from all seven continents) and affiliated these enzymes with a broader-than-expected distribution of taxonomic lineages. Our results indicate that PHA depolymerases are globally distributed and occur in most ecosystems. However, PHA depolymerases were not universally detected in metagenomic datasets (*e*.*g*., less than a third of the 1,492 aquatic metagenomes screened had predicted hits) or occurred with low frequency (Supp Fig 1). PHAmcl depolymerases were particularly poorly represented in the metagenomic dataset, with only 3% of aquatic metagenomes and no solid waste metagenomes harbouring predicted enzymes. There are very few reports of isolated PHAmcl degraders in the literature^22^ and our screen supports a sparse environmental distribution of these organisms. We allowed a permissive size cut-off threshold of 0.5Mb of assembled sequence data to retain even small datasets. Notably, the majority of thermal spring and volcanic metagenomes were small (81/121 and 4/5 metagenomes were <10Mb respectively), as well as a significant fraction of sediment, freshwater and activated sludge (212/442, 213/481 and 60/140 were <10Mb respectively). Since we were less likely to identify enzymes in smaller metagenomes, we performed normalization by dataset size to calculate PHA depolymerase frequency. We note that PHA depolymerases may be present in a greater proportion of environments than our screen predict. We explored the relationship between metagenome size and the frequency of predicted PHA depolymerases and did not find a strong linear relationship (R^2^= 0.0282) (Supp Fig 4). Predicted PHA depolymerases were placed in ML trees anchored with reference enzymes (Supp Fig 2). Based on tree placement, only a handful of predicted enzymes were closely related to biomarker enzymes; the majority of predicted enzymes formed more distantly related clades. If functional, these enzymes significantly expand the known sequence diversity of PHA depolymerases.

PHA depolymerase subtypes were predicted at different frequencies across ecosystems. PHAsclT1 depolymerases were the most prevalent followed by PHAsclT2 enzymes, with PHAmcl depolymerases having substantially lower frequencies across all ecosystems. Wastewater metagenomes, particularly those from activated sludge and nutrient removal systems, had the highest frequency of predicted PHA depolymerases and may be ecological hotspots for these enzymes. Interestingly, wastewater has been explored as a promising carbon source for PHA production^60^. With the predicted presence of PHA degraders in the majority of wastewater systems, there is the possibility of engineering a circular system where wastewater acts as both a feedstock and a waste processing component in the bioplastic lifecycle. Notably, a high frequency of PHAsclT1 depolymerases were detected in terrestrial metagenomes, with soil metagenomes harboring a huge number of novel predicted T1 enzymes, though at average or low relative frequencies.

Prior to our screen, bacterial ePHA depolymerases were characterized from the Proteobacteria, Actinobacteriota, Bdellovibrionota (formerly within the Deltaproteobacteria), and Firmicutes, as well as one fungal phylum, the Ascomycetes. Our screen infers PHA depolymerases occur in 23 additional bacterial phyla and 2 archaeal phyla. An LCA analysis on unbinned predicted PHA-depolymerase encoding scaffolds associated this predicted function with 28 different phylum-level lineages, 10 of which were also identified in our MAG screen. MAGs allowed us to more confidently taxonomically place putative PHA depolymerase hosts, compare enzyme frequencies between lineages, and place predicted enzymes within a genomic context whereas our work with metagenomic scaffolds provided a larger dataset originating from a greater number of environmental samples. Together, our analyses affiliated predicted PHA depolymerases predominantly with the Proteobacteria, Bdellovibrionota, Myxococcota, Methylomirabilota, Actinobacteriota, and Chloroflexota (Fig 3, Supp Fig 3). Many Proteobacterial PHA degraders have been identified through culture-based studies, particularly from soils^13^ and many of the PHA depolymerases used to seed our screen were from this phylum (Supp Data 1 and references within). For this reason, we cannot be certain if this phylum is genuinely enriched for this function compared to other phyla, meaning efforts to characterize distantly related enzymes from non-Proteobacterial lineages may confirm a broader phylogenetic distribution for this enzyme family.

A small number of predicted PHAscl depolymerase-encoding scaffolds identified from metagenomes were classified as archaeal, from the Halobacteriota, and one predicted PHAmcl depolymerase-encoding scaffold was classified as Thermoplasmatota. A number of distantly-related genes were identified in archaeal MAGs, but none passed our curation steps. These archaeal sequences may represent true PHA depolymerases that are distantly related from reference enzymes and thus may be interesting targets for future characterization (Supp Data 6).

Reference ePHAmcl depolymerases originated from the Bdellovibrionota (formerly a lineage within the Deltaproteobacteria), Gammaproteobacteria, and Actinobacteriota. We identified PHAmcl depolymerase-encoding MAGs from the Proteobacteria, Firmicutes and Myxococcota (formerly the Deltaproteobacteria), with the highest overall frequency occurring in the Firmicutes. More phylum-level diversity was found in the LCA analyses of PHAmcl depolymerase-encoding scaffolds, but interestingly no Firmicutes were identified. Most scaffolds were affiliated with the Proteobacteria, Myxococcota, and Actinobacteriota, but many were unclassified. Intriguingly, a few ePHAmcl-encoding scaffolds were classified as Acidobacteriota, Deinococcota, and Dormibacterota, which are lineages that have not yet been associated with PHAmcl depolymerases. T2 reference enzymes originated from the Proteobacteria (most), a fungal species (1) and Actinobacteriota (1). All MAG-encoded PHAsclT2 depolymerases occurred in members of the Proteobacteria (mainly the Burkholderiales) and often co-occurred with PHAsclT1 depolymerases. The restriction of predicted T2 enzymes to only a few lineages is an interesting result which was mirrored by our LCA analyses on T2-encoding scaffolds.

Two biochemically validated intracellular PHAsclT1 depolymerases (ACN29674.1 & BAE44206.1) are homologous to our reference extracellular PHAsclT1 depolymerases. To assess cellular localization of our predicted PHA depolymerases, we predicted signal peptides. The majority of MAG-encoded PHA depolymerases are expected to localize extracellularly. In contrast, fewer than half of the metagenome-derived PHA depolymerases had predicted signal peptides. We predict that a small fraction of identified PHA depolymerases may represent intracellular enzymes, but suspect that signal prediction algorithms may overlook valid signal peptides, particularly for organisms not well-represented in current databases.

We identified several hosts with multiple predicted PHA depolymerases, which may indicate metabolic flexibility on PHA substrates. We consider these organisms and their PHA depolymerases important targets for future characterization. Most notably, we identified a *Herminiimonas* species from the Burkholderiales which encoded a complement of nine predicted PHA depolymerases, suggesting it may be an efficient and/or highly versatile PHA degrader. We also identified a scaffold encoding two PHAmcl enzymes that was classified as from a member of the Myxococcota, indicating this family may be enriched for this rarer enzyme subtype.

Biodegradable plastics like PHA offer a promising solution to the devastating environmental effects caused by conventional plastics, however it is important to understand the potential impacts of adding a novel substrate into the environment. Our screen identified low frequencies or non-detection of PHA depolymerases in many environments. The presence of PHA waste may enrich low-abundance community members with PHA degradation capacity and significantly alter microbial community structure. In a controlled trial, biofilms associated with PHAs were dominated by sulfate-reducing microorganisms in comparison to polyethylene terephthalate controls^61^. The potential for low abundance organisms to respond to PHAs in an environment may allow more robust PHA degradation than our screen predicts, but also may have unexpected impacts on microbial community dynamics and biogeochemical cycling.

In conclusion, our work presents a carefully curated set of predicted PHA depolymerases to conservatively catalogue the existing environmental sequence diversity and the associated taxonomic affiliations of this enzyme family. We dramatically expand the inventory of PHA depolymerases available to target for future biochemical characterization, and affiliate many of these enzymes with novel taxonomic lineages for this function. We also provide an overview of the occurrence and frequency of PHA depolymerases detected in the environment. Future research should employ a combination of metagenomic predictions, possibly leveraging hidden markov models (HMMs) for broader diversity capture, burial degradation trials with functional profiling, and biochemical validation of predicted enzymes (particularly for enzymes distantly related to biomarker enzymes) in targeted environments to confirm predictions developed here.

## Data Availability

All predicted PHA depolymerase faa files, MSAs, and tree files are available at: https://github.com/vrviljak/-The-phylogenetic-and-global-distribution-of-polyhydroxyalkanoate-bioplastic-degrading-genes.git. All trees visualized with iTOL for this project can be viewed at: https://itol.embl.de/shared/2DHP8UFMQVeYO

## Acknowledgements

We would like to thank Angus Hilts for help with technical infrastructure. We would also like to extend our gratitude to the scientific community sequencing environmental metagenomes for their part in developing global data repositories. LAH is supported by a Tier II Canada Research Chair, and this work was funded by an NSERC Discovery grant (2016-03686) to LAH. A significant proportion of the datasets used were derived from grant support to Jill Banfield under DOE Office of Science BER grant DOE-SC10010566 and National Science Foundation grant CZP EAR-1331940; and to James Tiedje under DOE Office of Science BER DE-FC02-07ER64494.

## Supplementary Figure Legends

**Supplementary Figure 1:** The distribution of predicted PHA depolymerases showing geocoordinates of metagenomes containing predicted PHA depolymerases. Metagenomes are colored by the frequency of predicted PHA depolymerases in hits/Gb. For metagenomes sharing geocoordinates, the metagenome with the highest frequency of predicted enzymes is shown.

**Supplementary Figure 2:** Environmental origin and phylogenetic relationships of the 13,869 predicted PHA depolymerases identified in 1,295 metagenomes. Colored ring indicates the environmental origin of the metagenome encoding each predicted PHA depolymerase. Biomarker reference enzymes are indicated with black arrows. Protein trees were inferred with FastTree and show the sequence diversity of A) 222 predicted PHAmcl depolymerases B) 10,245 predicted ePHAsclT1 depolymerases and C) 3,402 predicted PHAsclT2 depolymerases, each with biochemically confirmed biomarker proteins included as references. Tree scale bar indicates number of substitutions per site.

**Supplementary Figure 3:** The phylogenetic affiliation of predicted PHA depolymerase enzymes on scaffolds >2,000 bps in length from 1,295 metagenomes using an LCA approach (as implemented in MEGAN). Shown is the relative abundance of various phyla and classes (for Proteobacteria only) of predicted PHA depolymerases for each enzyme subtype. “Unassigned” proteins could not be not classified at the phylum-level and all phyla that comprised less than 1% of proteins were combined into the “other” category. Number of scaffolds (n) for each enzyme category is provided.

**Supplementary Figure 4:** Plot showing the relationship between assembled metagenome size (Gb) and predicted PHA depolymerase frequency (hits/Gb). Plot shows a very weak positive linear relationship (R^2^ value = 0.0282) when Y-intercept was set to 0. Metagenomes are colored by environment of origin. Area captured in gray box is expanded in figure inset.

## Supplementary Data Files

All supplementary data files can be accessed through the Open Science Framework at doi: 10.17605/OSF.IO/VQTM6. Supplemental Data 5 is only available via OSF due to its large size. **Supplementary Data 1:** Supp_Data_1_PHA_depolymerases_references.xlsx

**Supplementary Data 2:** Supp_Data_2_PHA_depolymerases_from_global_metagenomes.xlsx

**Supplementary Data 3:** Supp_Data_3_PHA_depolymerase_MAGs.xlsx

**Supplementary Data 4:** Supp_Data_4_Detailed_PHA_depol_curation_methods.txt

**Supplementary Data 5:**

Supp_Data_5_BLAST_output_of_predicted_PHA_depolymerase_summary.xlsx can be accessed via the Open Science Framework at doi: 10.17605/OSF.IO/VQTM6

**Supplementary Data 6:** Supp_Data_6_unique_archaeal_IMG_MAG_hits_e10.xlsx

## Notes

### Competing Interest Statement

The authors have declared no competing interest.

### Summary of Updates

Revised article following two rounds of peer review.

